# Histone H3K4 methyltransferase DcATX1 promotes ethylene induced petal senescence in carnation

**DOI:** 10.1101/2022.01.12.476045

**Authors:** Shan Feng, Ruiming Wang, Hualiang Tan, Linlin Zhong, Yunjiang Cheng, Manzhu Bao, Hong Qiao, Fan Zhang

**Affiliations:** Key Laboratory of Horticultural Plant Biology, Ministry of Education, College of Horticulture and Forestry Sciences, Huazhong Agricultural University, Wuhan 430070, China; National R&D Center for Citrus Postharvest Technology, College of Horticulture and Forestry Sciences, Huazhong Agricultural University, Wuhan 430070, China; Hubei Hongshan Laboratory, Wuhan 430070, China; Key Laboratory of Urban Agriculture in Central China (pilot run), Ministry of Agriculture, College of Horticulture and Forestry Sciences, Huazhong Agricultural University, Wuhan 430070, China; Institute for Cellular and Molecular Biology, The University of Texas at Austin, Austin, Texas 78712, USA; Department of Molecular Biosciences, The University of Texas at Austin, Austin, Texas 78712, USA

## Abstract

Petal senescence is controlled by a complex regulatory network. Epigenetic regulation like histone modification influences chromatin state and gene expression. However, involvement of histone methylation in regulating petal senescence is still largely unknown. Here, we found that the trimethylation of histone H3 at Lysine 4 (H3K4me3) is increased during the ethylene induced petal senescence in carnation (*Dianthus caryophyllus* L.). The H3K4me3 levels are positively associated with the expression of transcription factor *DcWRKY75*, ethylene biosynthetic genes *DcACS1* and *DcACO1*, and senescence associated genes (SAGs) *DcSAG12* and *DcSAG29*. Further, we identified that carnation DcATX1 (ARABIDOPSIS HOMOLOG OF TRITHORAX1) encodes a histone lysine methyltransferase which can methylate H3K4. Knockdown of *DcATX1* delays ethylene induced petal senescence in carnation, which is associated with the downregulated expression of *DcWRKY75*, *DcACO1* and *DcSAG12*. While overexpression of *DcATX1* exhibits the opposite effects. DcATX1 promotes the transcription of *DcWRKY75*, *DcACO1* and *DcSAG12* by targeting to their promoters to elevate the H3K4me3 levels. Overall, our results demonstrate that DcATX1 is a H3K4 methyltransferase that promotes the expression of *DcWRKY75*, *DcACO1* and *DcSAG12* by regulating H3K4me3 levels, thereby accelerating ethylene induced petal senescence in carnation. This study further indicates that epigenetic regulation is important for plant senescence process.

**One sentence summary:** A histone methyltransferase promotes ethylene induced petal senescence in cut flower

## Introduction

Fresh flowers are essential for plant reproduction and human spiritual life, they are also important resources of spices, teas and pigments. The process of flower senescence, especially petal senescence largely determines the ornamental and economic value of a flower. Elucidating the molecular mechanisms of petal senescence not only contributes to the improving of the postharvest longevity of cut flowers but also to the entire floral industry (Ma et al., 2018).

Petal senescence is the final step of floral development that is highly regulated by a complex network of both endogenous hormones and exogenous factors (Ma et al., 2018). Among them, ethylene is considered to be the major hormone which regulates the senescence process of ethylene sensitive flowers (Ma et al., 2018). Carnation (*Dianthus caryophyllus* L.) is one of the most important and universally used ornamental cut flowers worldwide which is highly sensitive to ethylene, so it is generally served as a model plant for studying the mechanism of ethylene induced petal senescence (Yagi et al., 2014; Ma et al., 2018).

Ethylene, a gaseous plant hormone, is essential for multiple developmental and physiological processes in plants and responses to biotic and abiotic stresses (Merchante et al., 2013; Ju and Chang, 2015). The ethylene response is initiated by ethylene biosynthesis and signaling transduction (Yang and Hoffman, 1984; Merchante et al., 2013; Ju and Chang, 2015). In ethylene biosynthesis pathway, the 1-aminocyclopropane-1-carboxylic acid (ACC) synthase (ACS) and ACC oxidase (ACO) play key roles (Yang and Hoffman, 1984). In addition, the ethylene signaling transduction pathway has been well characterized in model plant *Arabidopsis* (Wang and Qiao, 2019; Binder, 2020; Wang et al., 2020; Zhao et al., 2021). In brief, ethylene is perceived by receptors (Chang et al., 1993; Hua et al., 1995; Hua and Meyerowitz, 1998; Hua et al., 1998). Then the receptors may undergo conformational changes and fail to activate CONSTITUTIVE TRIPLE RESPONSE1 (CTR1) (Kieber et al., 1993), a kinase of ETHYLENE INSENSITIVE2 (EIN2) (Alonso et al., 1999; Ju et al., 2012).

So, the C-terminal end of EIN2 (EIN2-C) will be cleaved, and then translocated into cytoplasmic processing body (P-body) and nucleus (Ju et al., 2012; Qiao et al., 2012; Wen et al., 2012; Li et al., 2015; Merchante et al., 2015). In P-body, EIN2-C will repress the translation of EIN3-BINDING F BOX PROTEIN1 (EBF1) and EBF2 to stabilize two core transcription factors EIN3 and EIN3-LIKE1 (EIL1) (Chao et al., 1997; Guo and Ecker, 2003; Potuschak et al., 2003; Gagne et al., 2004; Li et al., 2015; Merchante et al., 2015); In nucleus, EIN2-C will transduce the signal to EIN3 and EIL1, leading to the transcriptional activation of ethylene response genes (Solano et al., 1998; Chang et al., 2013). Recent studies showed that under ethylene treatment, EIN2 interacts with a histone binding protein EIN2 NUCLEAR ASSOCIATED PROTEIN1 (ENAP1) to elevate the acetylation levels of histone H3 at Lysine 14 and 23 (H3K14Ac and H3K23Ac), further initiates the transcriptional activation of downstream genes (Zhang et al., 2016; Wang et al., 2017; Zhang et al., 2017; Zhang et al., 2018; Wang et al., 2021). This links the ethylene signaling transduction pathway with epigenetic regulation like histone modification (Wang and Qiao, 2019, 2020; Wang et al., 2020).

According to the functional validation and gene expression patterns analysis, ethylene biosynthesis and signaling transduction genes such as *DcACS1*, *DcACO1*, *DcEIN2*, *DcEIL3* and *DcEBF1* may play vital roles in ethylene induced petal senescence in carnation (Park et al., 1992; Woodson et al., 1992; Savin et al., 1995; tenHave and Woltering, 1997; Jones and Woodson, 1999; Waki et al., 2001; Shibuya et al., 2002; Iordachescu and Verlinden, 2005; Fu et al., 2011; Fu et al., 2011). Recently, a study indicated that transcription factors DcEIL3 and DcWRKY75 play key positive roles in ethylene induced petal senescence in carnation (Xu et al., 2021). However, the regulation networks and mechanisms, especially the epigenetic regulation mechanisms remain largely unidentified.

Histone modification such as histone methylation has been implicated to play critical roles in senescence process in plants, mainly in leaf senescence (Ay et al., 2014; Woo et al., 2019; Ostrowska-Mazurek et al., 2020). A previous study revealed that activation of *WRKY53*, a key regulator of leaf senescence, correlated with a significant increase in H3K4 methylation level (Ay et al., 2009). Overexpression of *SU(VAR)3-9 HOMOLOG2* (*SUVH2*), a histone methyltransferase, significantly decreases H3K4 methylation level but increases H3K27 methylation levels at 5’ end and coding regions of *WRKY53* (Ay et al., 2009). Further, genome-wide analysis showed that the expression of SAGs is positively associated with the H3K4me3 levels but negatively associated with the H3K27me3 levels during leaf senescence (Brusslan et al., 2012; Brusslan et al., 2015). JUMONJI DOMAIN-CONTAINING PROTEIN16 (JMJ16), a histone H3K4 demethylase, was found to repress leaf senescence in *Arabidopsis* by targeting to the promoters of *WRKY53* and *SAG201* (Liu et al., 2019); In addition, RELATIVE OF EARLY FLOWERING6 (REF6), a H3K27me3 demethylase, was found to promote leaf senescence through directly activating the major senescence regulatory and functional genes in *Arabidopsis* (Wang et al., 2019), further confirming the importance of histone methylation in leaf senescence.

Although H3K4 methylation has been shown to be important for leaf senescence, whether and how it participates in ethylene induced petal senescence in carnation is still largely unexplored. In this study, we found that during the ethylene induced carnation petal senescence, the H3K4me3 levels on the promoter regions of *DcWRKY75*, *DcACS1*, *DcACO1*, *DcSAG12* and *DcSAG29* were increased. We then showed that DcATX1 is a H3K4 methyltransferase that promotes the ethylene induced carnation petal senescence process. Finally, we demonstrated that DcATX1 binds to the promoters of *DcWRKY75*, *DcACO1* and *DcSAG12* to increase the H3K4me3 levels, thereby inducing the expression of *DcWRKY75*, *DcACO1* and *DcSAG12*. Together, our research revealed that DcATX1 is a H3K4 methyltransferase that regulates the transcription of *DcWRKY75*, *DcACO1* and *DcSAG12* by modulating H3K4me3 levels to promote the ethylene induced petal senescence in carnation.

## Results

### H3K4me3 levels are elevated during ethylene induced petal senescence in carnation

Our previous study has shown that the Gene Ontology (GO) analysis of 2973 Cluster 8 genes in ethylene treated carnation petal transcriptome were summarized in the stress, ethylene and aging pathway, which indicated that these genes might play key roles in ethylene induced petal senescence in carnation (Xu et al., 2021). We also found that the GO term of regulation of histone modification like histone methylation and histone acetylation exist in this cluster (Xu et al., 2021), which suggested that histone modification may also play important roles in ethylene induced petal senescence in carnation.

To test whether histone methylation like H3K4me3 involved in carnation petal senescence, we firstly detected the H3K4me3 modification level in this process. Previous study has shown that the carnation flower opening and senescence process can be divided into six stages: bud stage (BS), half bloom stage (HBS), full bloom stage (FBS), beginning of wilting stage (BWS), half wilting stage (HWS) and complete wilting stage (CWS) (Fig. 1a) (Kong et al., 2017). We collected petals at different stages and examined the total H3K4me3 levels by western blot. H3K4me3 levels were obviously increased at the BWS stage and were maintained at high levels during the senescence (Fig. 1b), showing that the H3K4me3 levels are indeed elevated during the petal senescence in carnation.

**Figure 1.**
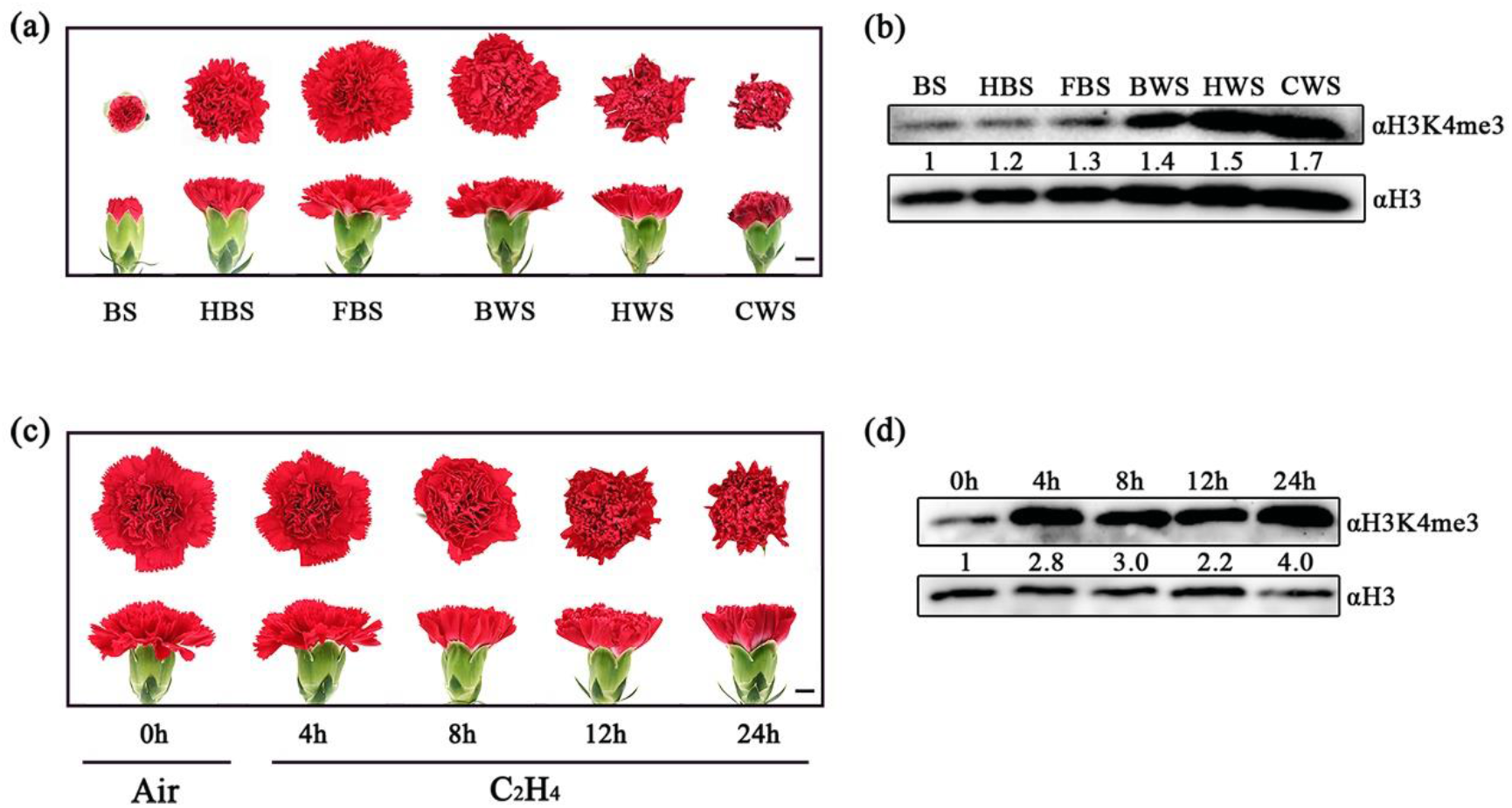
Ethylene induces carnation petal senescence and elevates H3K4me3 level. **(a)** Phenotype of carnation flowers during opening and senescence process. Scale bar = 1 cm. **(b)** Western blot analysis of the H3K4me3 modification level of carnation petal during opening and senescence process. An anti-H3K4me3 antibody was used for modification detection, and an anti-H3 antibody was used for detection of loading control. Relative amounts of H3K4me3 modification normalized to H3 are shown below. **(c)** Phenotype of FBS carnation flowers treated with ethylene by different times. Scale bar = 1 cm. **(d)** Western blot analysis of the H3K4me3 modification level of carnation petal under ethylene treatment. An anti-H3K4me3 antibody was used for modification detection, and an anti-H3 antibody was used for detection of loading control. Relative amounts of H3K4me3 modification normalized to H3 are shown below.

To further examine whether ethylene promotes H3K4me3 accumulation during the petal senescence process in carnation, we firstly examined the phenotype of carnation flower senescence at FBS stage treated with 10 ppm ethylene for different times (0 hour (h), 4h, 8h, 12h and 24h). We found that 8h ethylene treatment can cause a visible petal wilting phenotype, and the phenotype was severer with 24h ethylene treatment (Fig. 1c). This phenotype change is accordance with our previous study (Xu et al., 2021). We then conducted western blot to examine the H3K4me3 levels in those flower petals, and we found that the H3K4me3 levels were obviously accumulated in the petals with 4 hours of ethylene treatment, and the levels were maintained thereafter (Fig. 1d), indicating that H3K4me3 may play important roles in carnation petal senescence.

These data suggested that H3K4me3 levels are elevated during ethylene induced petal senescence in carnation.

### H3K4me3 modifications are positively associated with expression levels of *DcWRKY75*, *DcACS1*, *DcACO1*, *DcSAG12* and *DcSAG29* in carnation

To detect whether the increased H3K4me3 level influence gene expression, we selected several senescence associated key genes to verify our hypothesis.

Previous study indicated that a strong correlation between changes in the H3K4me3 mark and gene expression of *WRKY75* was observed during leaf senescence in *Arabidopsis* (Brusslan et al., 2015). Since *DcWRKY75*, a homolog gene of *WRKY75* in *Arabidopsis*, is a key positive regulator in ethylene induced petal senescence in carnation and belongs to Cluster 8 (Xu et al., 2021), we firstly detect the expression level of *DcWRKY75* and H3K4me3 modification in the promoter region of *DcWRKY75*. Quantitative Reverse Transcription PCR (RT-qPCR) assay indicated that *DcWRKY75* is quickly induced by ethylene treatment (Fig. 2a), which is accordance with our previous study (Xu et al., 2021). Chromatin Immunoprecipitation (ChIP)-qPCR assay exhibited that H3K4me3 levels in the different promoter regions of *DcWRKY75* were highly enriched by ethylene treatment (Fig. 2b). This means that ethylene indeed can increase the H3K4me3 level in the promoter region of *DcWRKY75* to enhance its expression.

**Figure 2.**
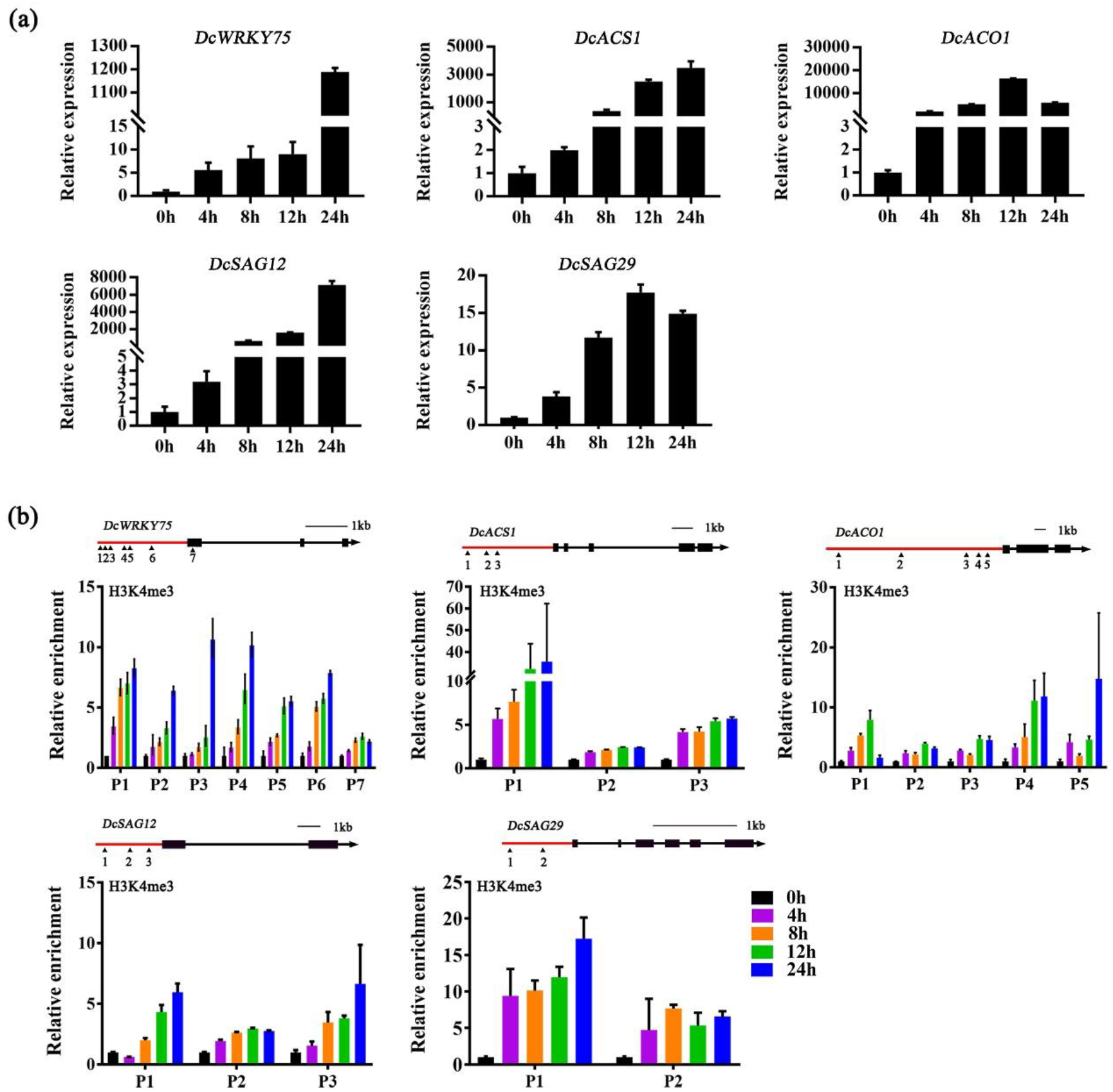
Relative expression of *DcWRKY75*, *DcACS1*, *DcACO1*, *DcSAG12* and *DcSAG29* and H3K4me3 enrichments in the promoter regions of these genes in carnation petal under ethylene treatment. **(a)** Relative expression of *DcWRKY75*, *DcACS1*, *DcACO1*, *DcSAG12* and *DcSAG29* in carnation petal under ethylene treatment. **(b)** ChIP-qPCR analysis of H3K4me3 enrichment at promoters of *DcWRKY75*, *DcACS1*, *DcACO1*, *DcSAG12* and *DcSAG29* in carnation petal under ethylene treatment. Schematic structure of genomic sequences of *DcWRKY75*, *DcACS1*, *DcACO1*, *DcSAG12* and *DcSAG29* were shown. Arabic numbers indicate the sites at *DcWRKY75*, *DcACS1*, *DcACO1*, *DcSAG12* and *DcSAG29* locus used for ChIP-qPCR analysis. Red lines represent promoter regions, black bars represent exons, and black lines represent intron regions.

Since ethylene biosynthetic genes *DcACS1* and *DcACO1* and senescence associated genes *DcSAG12* and *DcSAG29* also play critical roles in ethylene induced petal senescence in carnation (Xu et al., 2021), we examined the expression levels of these genes by ethylene treatment. RT-qPCR assay indicated that *DcACS1*, *DcACO1*, *DcSAG12* and *DcSAG29* are also obviously upregulated by ethylene treatment (Fig. 2a), which is also accordance with our previous study (Xu et al., 2021). ChIP-qPCR assay exhibited that H3K4me3 levels in the different promoter regions of *DcACS1*, *DcACO1*, *DcSAG12* and *DcSAG29* were also highly enriched by ethylene treatment (Fig. 2b). These mean that ethylene indeed can also increase the H3K4me3 levels in the promoter region of *DcACS1*, *DcACO1*, *DcSAG12* and *DcSAG29* to enhance their expression.

Overall, these data showed that H3K4me3 levels are positively associated with expression levels of *DcWRKY75*, *DcACS1*, *DcACO1*, *DcSAG12* and *DcSAG29*, which indicated that H3K4me3 is involved in in ethylene induced petal senescence process in carnation.

### DcATX1 is a potential H3K4 methyltransferase in carnation

In higher plants, histone methylation levels are dynamically regulated by histone methyltransferases and histone demethylases, which are mainly conducted by SET DOMAIN GROUP (SDG) proteins and JMJ proteins (Cheng et al., 2020). To illustrate how ethylene promotes H3K4me3 accumulation during the petal senescence process of carnation, we firstly analyze the SDG proteins which have potential H3K4 methyltransferase activity in carnation.

In model plant *Arabidopsis*, ATX1/SDG27, ATX2/SDG30, ATX3/SDG14, ATX4/SDG16, ATX5/SDG29, ARABIDOPSIS TRITHORAX-RELATED3 (ATXR3)/SDG2, ATXR7/SDG25, ABSENT, SMALL, OR HOMEOTIC DISCS1 HOMOLOG1 (ASHH1)/SDG26, ASHH2/SDG8 and ASH1-RELATED3 (ASHR3)/SDG4 have been reported to regulate the H3K4 methylation level (Cartagena et al., 2008; Saleh et al., 2008; Cazzonelli et al., 2009; Tamada et al., 2009; Berr et al., 2010; Guo et al., 2010; Berr et al., 2015; Chen et al., 2017; Cheng et al., 2020). eFP browser (http://bar.utoronto.ca/efp/cgi-bin/efpWeb.cgi) show that among these genes, only *ATX1* exhibits an increased expression in petals and stamen at flower stage 15 by comparing with petals and stamen at flower stage 12 (Fig. S1), which means that ATX1 homolog proteins may participate in petal senescence process in *Arabidopsis* and other plants like in carnation.

By searching the homolog proteins in Carnation Genome Database (http://carnation.kazusa.or.jp/) use the above SDG proteins as query, we totally got 12 homolog proteins in carnation genome: DcATX1 (Dca16956), DcATX3 (Dca38), DcATX4 (Dca52395), DcATXR3-1 (Dca19660), DcATXR3-2 (Dca19661), DcATXR3-3 (Dca28546), DcATXR3-4 (Dca28547), DcATXR7 (Dca30048/Dca34598), DcASHH1-1 (Dca3622), DcASHH1-2 (Dca46518) and DcASHH2 (Dca26204) (Fig. S2a; Data S1) (Yagi et al., 2014). Based on our previously constructed ethylene treated carnation petal transcriptome (Xu et al., 2021), the heatmap showed that *DcATX1* exhibited an obvious down-regulated expression with ethylene treatment by comparing with other SDG genes (Fig. S2b; Data S1). RT-qPCR assay indeed indicated that the expression level of *DcATX1* showed a gradual decrease expression pattern during the ethylene induced petal senescence process (Fig. 3a).

**Figure 3.**
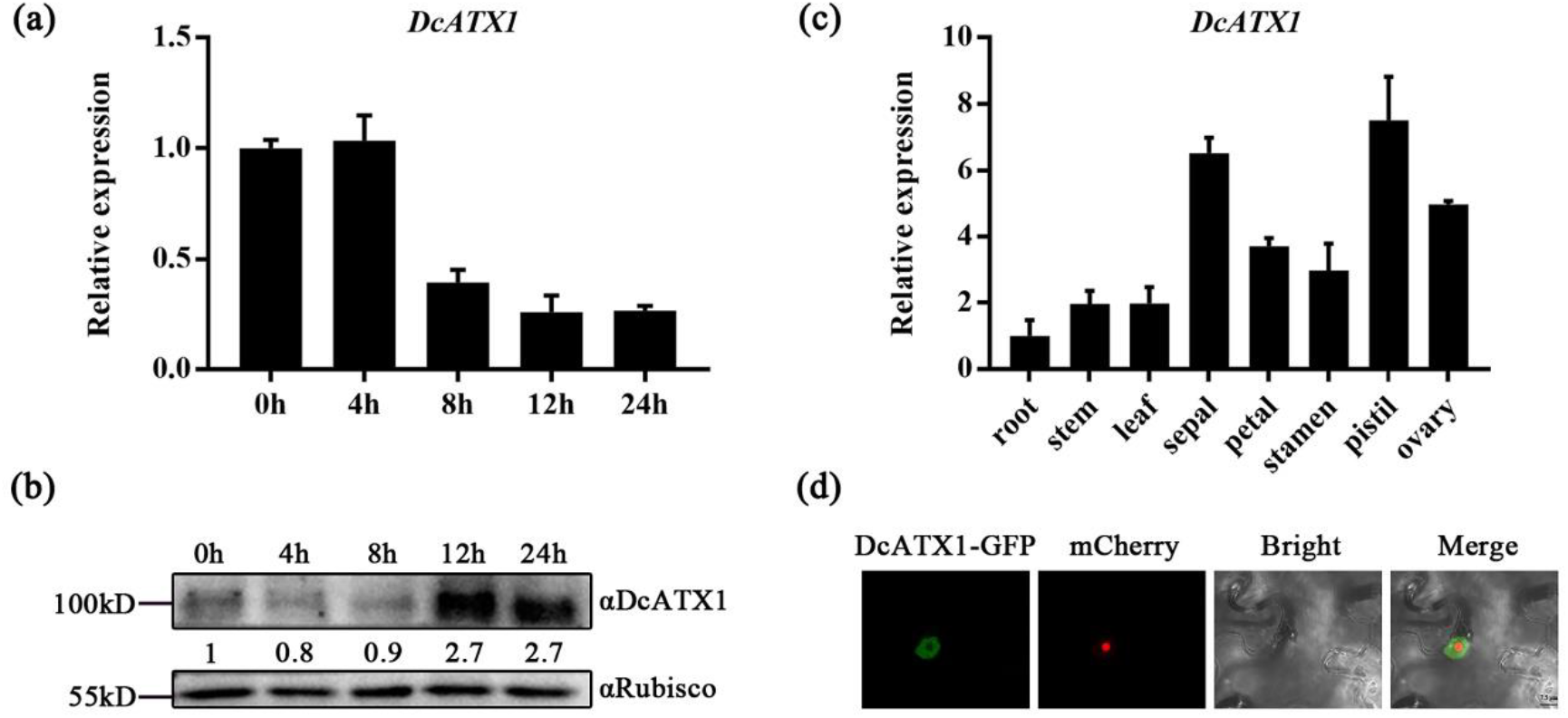
Identification of DcATX1 in carnation. **(a)** Relative expression of *DcATX1* in carnation petal under ethylene treatment. **(b)** DcATX1 protein level in carnation petal under ethylene treatment. Anti-Rubisco were used for detection of loading control. Relative amounts of DcATX1 levels normalized to Rubisco are shown below. ‘kD’ means ‘kilo Dalton’. **(c)** Relative expression of *DcATX1* in root, stem, leaf, sepal, petal, stamen, pistil and ovary of carnation. **(d)** Subcellular localization of DcATX1. Red fluorescent was a nucleoid maker. Scale bar = 7.5 µm.

Protein sequence alignment indicated that DcATX1 indeed had a typical SET (**<u>s</u>**uppressor of variegation, **<u>e</u>**nhancer of zeste and **<u>t</u>**rithorax) domain, which is critical for histone methyltransferase activity (Fig. S3). Further, we detected the protein level of DcATX1 using DcATX1 native antibody in carnation petal under ethylene treatment. To our surprise, the protein content of DcATX1 showed an obvious elevation by 12h ethylene treatment (Fig. 3b). This may due to the negative feedback regulation which indicated that DcATX1 might play important role in ethylene induced petal senescence in carnation, even though it showed an opposite expression pattern by comparing with *ATX1* in *Arabidopsis* (Fig. S1), so we selected it for further investigation.

We checked the tissue specific expression pattern of *DcATX1* and found that it showed a moderate expression level in petal (Fig. 3c). Further, we found that DcATX1 is localized in nucleus (Fig. 3d). These data indicate that DcATX1 is a potential H3K4 methyltransferase in ethylene induced petal senescence in carnation.

### Silencing of *DcATX1* delays ethylene induced petal senescence in carnation

To test the potential role of DcATX1 in petal senescence, we investigated its function by using virus-induced gene silencing (VIGS) technique (Cheng et al., 2018; Xu et al., 2021). We constructed a tobacco rattle virus vector (TRV-*DcATX1*) to specifically silence *DcATX1* in carnation plants and observed the phenotype. Compared with the TRV control plants, *DcATX1* exhibited reduced expression level in TRV-*DcATX1* silenced plants (Fig. 4c).

**Figure 4.**
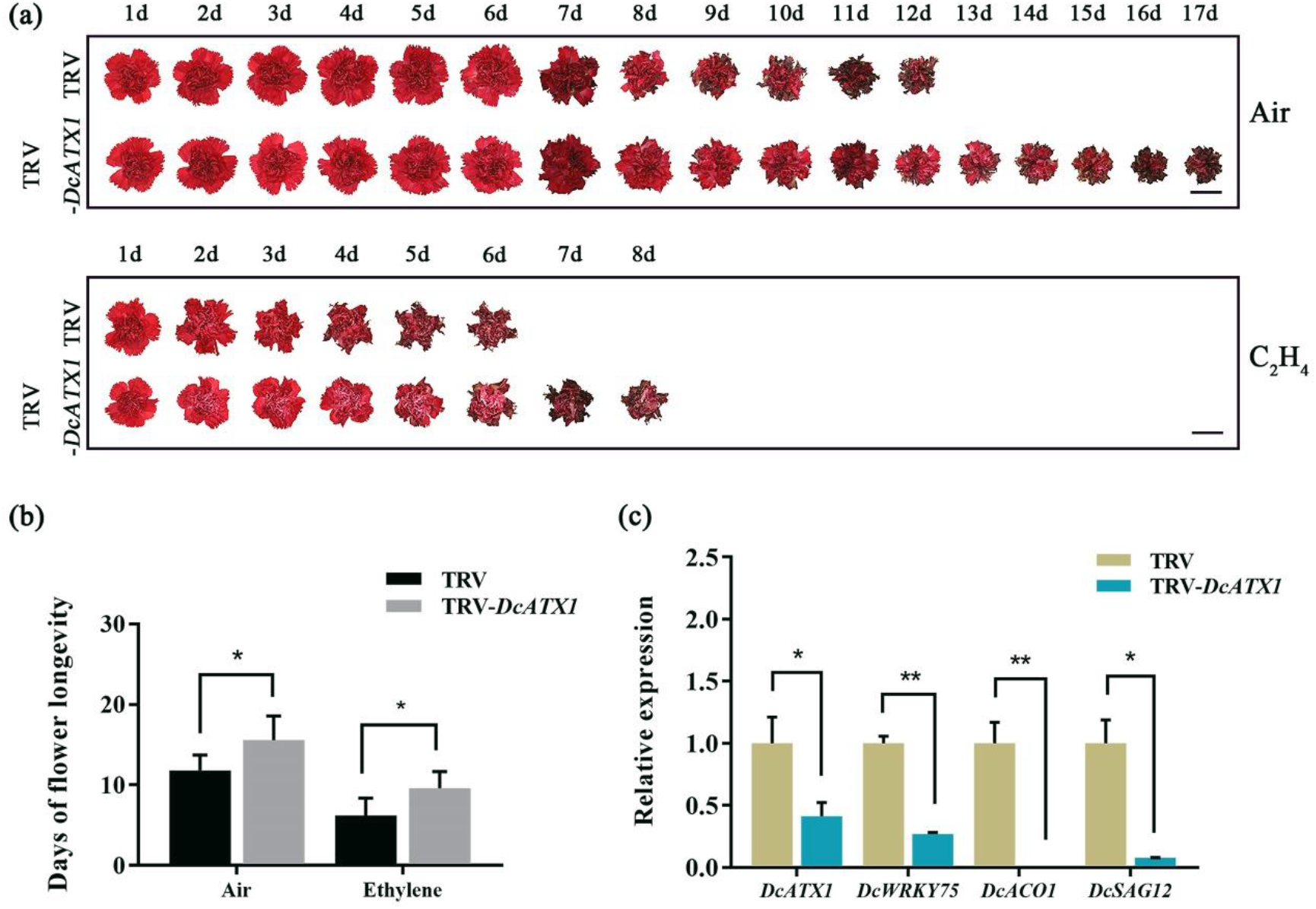
Silencing of *DcATX1* delays ethylene induced petal senescence in carnation. **(a)** Phenotype of carnation petal senescence of TRV control plants and TRV-*DcATX1* silenced plants without (Air) and with (C_2_H_4_) ethylene treatment, recorded daily. Scale bar = 5 cm. **(b)** The days of flower longevity of TRV control plants and TRV-*DcATX1* silenced plants without (Air) and with (C_2_H_4_) ethylene treatment. **(c)** Relative expression of *DcATX1*, *DcWRKY75*, *DcACO1* and *DcSAG12* in TRV control plants and TRV-*DcATX1* silenced plants.

Under air condition, the TRV-*DcATX1* silencing clearly delayed carnation petal senescence compared with in the TRV control plants, with flower longevity lasting 15.6±3.0days (d) in TRV-*DcATX1* silenced plants compared with 11.8±1.9d in the TRV control plants (Figs. 4a, b). After ethylene treatment, the petal senescence of both the TRV control plants and TRV-*DcATX1* silenced plants were accelerated, however, the vase life of the TRV-*DcATX1* silenced plants (9.6±2.1d) remained longer than that of the TRV control plants (6.2±2.2d) (Fig. 4a, b). RT-qPCR showed that the expression of *DcWRKY75*, *DcACO1* and *DcSAG12* were significantly decreased in TRV-*DcATX1* silenced plants compared with in the TRV control plants (Fig. 4c).

We also detected the potential function of DcATX1 in carnation petal discs. The effects of *DcATX1* silencing on senescence of petal discs were analyzed and the result showed that the expression of *DcATX1* was significantly lower in *DcATX1*-silenced petal discs than in the TRV control petal discs under ethylene treatment or not (Fig. S4b). In the TRV control, slight color fading occurred at 5d and almost half of the petal discs were discolored at 9d (Fig. S4a). In contrast, TRV-*DcATX1* silenced petal discs showed a delayed color fading phenotype, with only slight color fading at 9d (Fig. S4a). After ethylene treatment, color fading occurred at 5d and almost all the petal discs were discolored at 9d in the TRV control petal discs (Fig. S4a). In contrast, TRV-*DcATX1* silenced petal discs clearly exhibited a delayed senescence phenotype, with only slight color fading at 6d (Fig. S4a). In addition, the ion leakage rate showed a consistent trend with the expression level of *DcATX1* in different samples (Fig. S4c). RT-qPCR showed that the expression of *DcWRKY75* and *DcACO1* were significantly decreased in TRV-*DcATX1* silenced petal discs compared with in the TRV control petal discs no matter with or without ethylene treatment (Fig. S4d), but *DcSAG12* was significantly decreased in TRV-*DcATX1* silenced petal discs only in air condition (Fig. S4d).

These data indicated that silencing of *DcATX1* delays ethylene induced petal senescence in carnation.

### Overexpression of *DcATX1* accelerates ethylene induced petal senescence in carnation

To further investigate the potential function of DcATX1 in carnation petal senescence, we construct constitutively expressed *DcATX1* vector under the control of the cauliflower mosaic virus (CaMV) 35S promoter (*35S:DcATX1*) to overexpress *DcATX1* in carnation plants. Compared with the *35S* control plants, *DcATX1* exhibited an obvious overexpressed expression level in *35S:DcATX1* overexpression plants (Fig. 5c).

**Figure 5.**
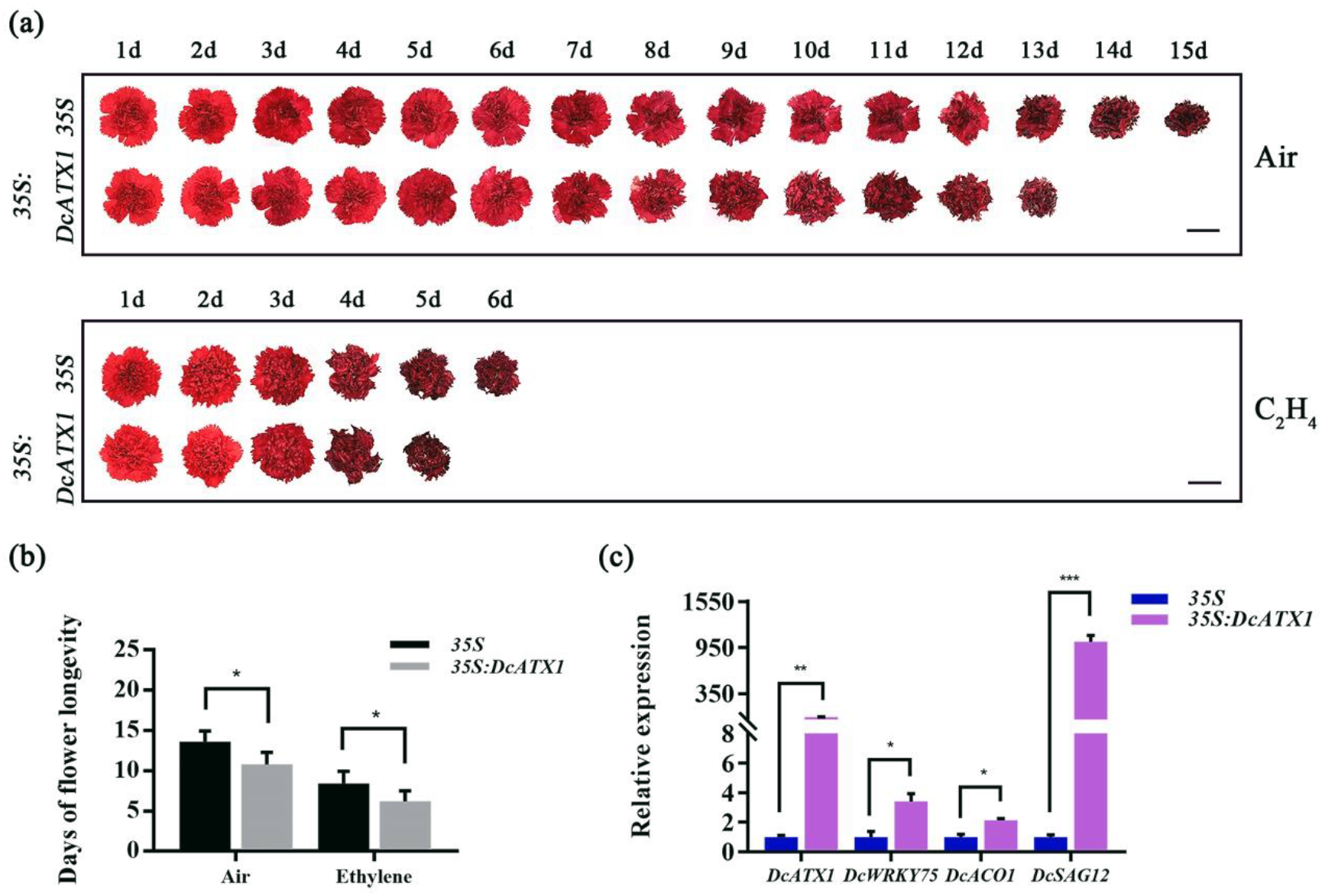
Overexpression of *DcATX1* accelerates carnation petal senescence. **(a)** Phenotype of carnation petal senescence of *35S* control plants and *35S:DcATX1* overexpression plants without (Air) and with (C_2_H_4_) ethylene treatment, recorded daily. Scale bar = 5 cm. **(b)** The days of flower longevity of *35S* control plants and *35S:DcATX1* overexpression plants without (Air) and with (C_2_H_4_) ethylene treatment. **(c)** Relative expression of *DcATX1*, *DcWRKY75*, *DcACO1* and *DcSAG12* in *35S* control plants and *35S:DcATX1* overexpression plants.

Under air condition, the *35S:DcATX1* overexpression plants showed a shorter life span than the *35S* control plants, with flower longevity lasting 10.8±1.5d in *35S:DcATX1* overexpression plants compared with 13.6±1.3d in the *35S* control plants (Fig. 5a, b). Under ethylene treatment, the flower longevity of the *35S:DcATX1* overexpression plants (6.2±1.3d) still remained shorter than that of the *35S* control plants (8.4±1.5d) (Fig. 5a, b). The RT-qPCR result showed that the expression of *DcWRKY75*, *DcACO1* and *DcSAG12* were significantly increased in *35S:DcATX1* overexpression plants compared with in the *35S* control plants (Fig. 5c).

We further tested the potential role of DcATX1 in carnation petal disc senescence and analyzed the effects of *DcATX1* overexpression on petal discs (Figs. S5b). Under air condition, slight color fading occurred at 12d and half of the petal discs were discolored at 15d in the *35S:DcATX1* overexpression petal discs (Fig. S5a). In contrast, *35S* control petal discs showed color fading only at 15d (Fig. S5a). After ethylene treatment, the *DcATX1* overexpressed petal discs were almost completely discolored at 15 d (Fig. S5a), whereas only half of the *35S* control petals discs showed color fading at 15d (Fig. S5a). In addition, the ion leakage rates were higher in *35S:DcATX1* overexpression petal discs at 3d under ethylene condition (Fig. S4c), and higher at 6d no matter with or without ethylene treatment (Fig. S4c). Accordance, the expression level of *DcWRKY75* were significantly increased in *35S:DcATX1* overexpression petal discs compared with in the *35S* control petal discs no matter with or without ethylene treatment (Fig. S4d), but *DcACO1* and *DcSAG12* were significantly increased in *35S:DcATX1* overexpression petal discs only under ethylene treatment (Fig. S4d). These data indicated that DcATX1 promotes ethylene induced petal senescence in carnation.

To further examine whether DcATX1 promotes petal senescence, we transformed the *DcATX1* overexpression vector (*35S:DcATX1*) into *Arabidopsis* Col-0 wild type (WT) and obtained three independent overexpression transgenic lines named *35S:DcATX1#1*, *35S:DcATX1#2* and *35S:DcATX1#3*. RT-qPCR result indicated that the expression level of *DcATX1* were strongly overexpressed in *35S:DcATX1* overexpression lines (Fig. S6a). The *35S:DcATX1* overexpression lines showed a significant accelerated phenotype in flower senescence and abscission compared with in WT (Fig. S6b). Consistently, RT-qPCR results showed a similar expression trends of senescence mark genes *AtSAG12* and *AtSAG29* with *DcATX1* in different lines (Fig. S6a). These results indicate that DcATX1 also promotes petal senescence in *Arabidopsis*.

### DcATX1 methylates H3K4 in carnation

To test whether DcATX1 regulates H3K4me3 modification, we examined the H3K4 methylation levels in TRV-*DcATX1* silenced plants and *35S:DcATX1* overexpression plants by western blot. We found that the H3K4me3 levels were significantly decreased in TRV-*DcATX1* silenced plants compared with in TRV control plants, and the H3K4me3 levels were significantly increased in *35S:DcATX1* overexpression plants compared with in *35S* control plants (Fig. 6a, b). Notably, the H3K4me2 levels were significantly increased in TRV-*DcATX1* silenced plants compare with that in TRV control plants, and significantly decreased in *35S:DcATX1* overexpression plants compared with that in *35S* control plants (Fig. 6a, b). However, no significant changes were detected in H3K4me1 levels (Fig. 6a, b). These results suggests that DcATX1 is a potential histone H3K4 methyltransferase that may methylate H3K4me2 into H3K4me3.

**Figure 6.**
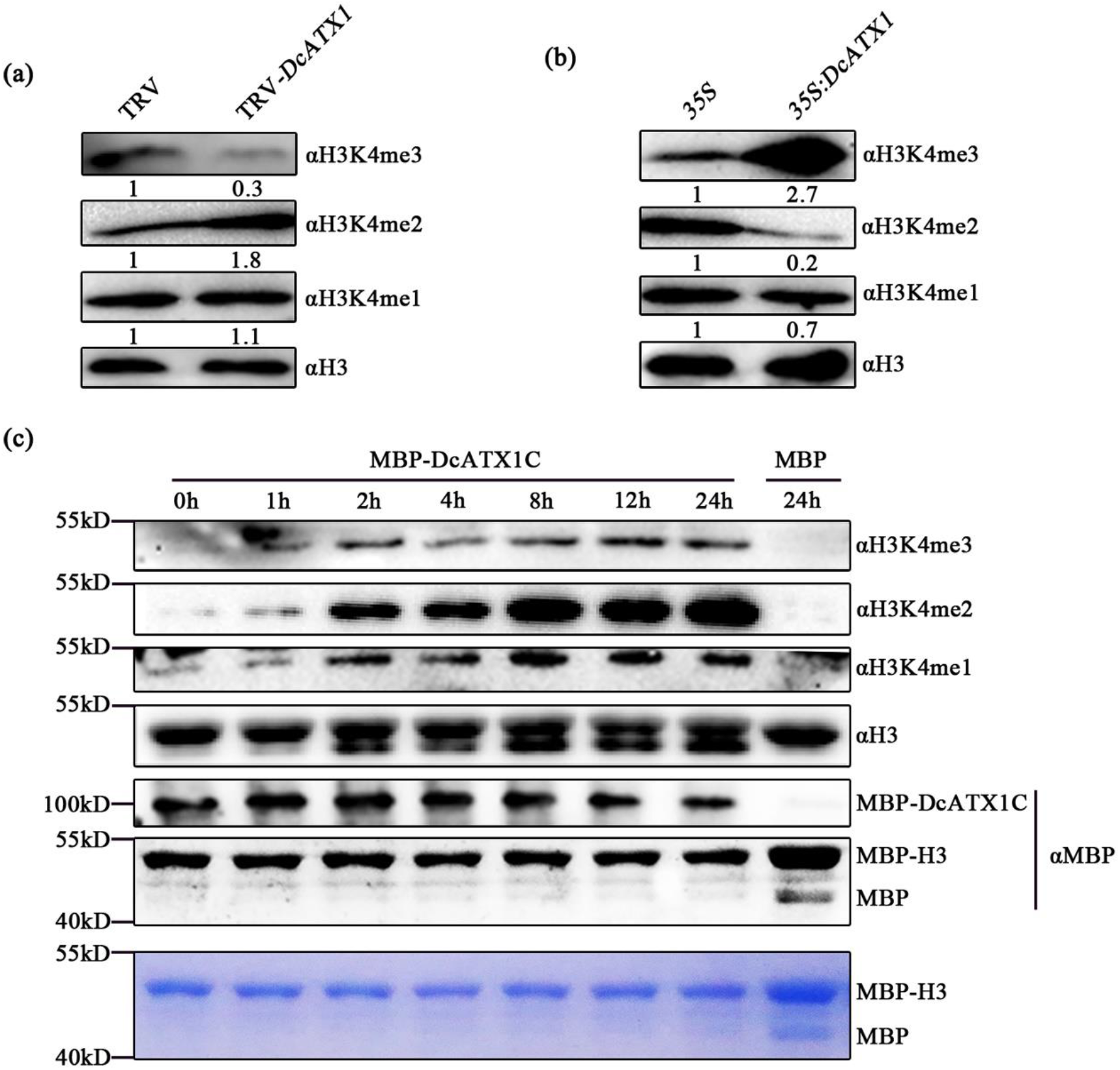
DcATX1 acts as a histone H3K4 methyltransferase. **(a, b)** H3K4 methylation levels in TRV-*DcATX1* silenced plants **(a)** and *35S:DcATX1* overexpression plants **(b)**. Protein extracted from carnation petals were analyzed by western blot using specific antibodies against H3K4me3, H3K4me2, H3K4me1 and H3 as indicated. Relative amounts of H3K4me3, H3K4me2, H3K4me1 modifications normalized to H3 are shown below. **(c)** DcATX1 has H3K4 methyltransferase activity. Recombinant *Arabidopsis* histone MBP-H3 were used as substrates and the mixture were incubated for 0-24h as indicated. The upper panels show western blots using specific antibodies against H3K4me3, H3K4me2, H3K4me1, H3 and MBP as indicated and the lower panels shows SDS-PAGE gel stained with coomassie brilliant blue R-250 for MBP-H3 and MBP. ‘kD’ means ‘kilo Dalton’.

In order to determine whether DcATX1 is indeed an active H3K4 methyltransferase, we did *in vitro* histone methyltransferase activity assay. We expressed and purified the C-terminal region of DcATX1 including the SET domain and fused to MBP tag (Fig. S3). The histone methyltransferase activity assay was performed using the fusion protein MBP-DcATX1C as enzyme source, a recombinant *Arabidopsis* histone MBP-H3 as the substrate and the S-adenosyl-L-methionine (SAM) as the methyl donor, MBP was used as the negative control. The reactions were incubated for various time periods from 1h to 24 h and the productions were analyzed by western blot using specific antibodies against H3K4me3, H3K4me2, H3K4me, H3 and MBP. H3K4me3 was detected within 1 h and progressively accumulated as the reaction time increase, well, no accumulation of H3K4me3 was detected in the MBP negative control even it was incubated for 24h (Fig. 6c). Same cases for the accumulation of H3K4me2 and H3K4me1 (Fig. 6c). This result indicated that DcATX1 can methylate recombinant *Arabidopsis* histone H3 rapidly which does not require any other proteins. Thus, we conclude that DcATX1 is a histone methyltransferase which is capable of catalyzing H3K4 methylation.

### DcATX1 promotes the transcription of *DcWRKY75*, *DcACO1* and *DcSAG12* by elevating H3K4me3 levels in their promoters

Since DcATX1 is a H3K4 methyltransferase and the expression levels of *DcWRKY75*, *DcACO1* and *DcSAG12* were decreased in TRV-*DcATX1* silenced plants and increased in *35S:DcATX1* overexpression plants, we want to detect whether DcATX1 regulates the H3K4me3 modification in the promoter regions of these genes.

We firstly did the ChIP-qPCR assays in TRV-*DcATX1* silenced plants and in TRV control plants with and without ethylene treatment. We found that in the promoter of *DcWRKY75*, H3K4me3 levels were obviously decreased at different regions no matter with or without ethylene treatment in TRV-*DcATX1* silenced plants, except for P3 region at air condition (Fig. 7a). For *DcACO1* promoter, H3K4me3 levels were obviously decreased at P2 and P5 regions under ethylene treatment in TRV-*DcATX1* silenced plants (Fig. 7a). For *DcSAG12* promoter, H3K4me3 levels were obviously decreased at P2 region no matter with or without ethylene treatment, but decreased at P1 region only in air condition in TRV-*DcATX1* silenced plants (Fig. 7a).

**Figure 7.**
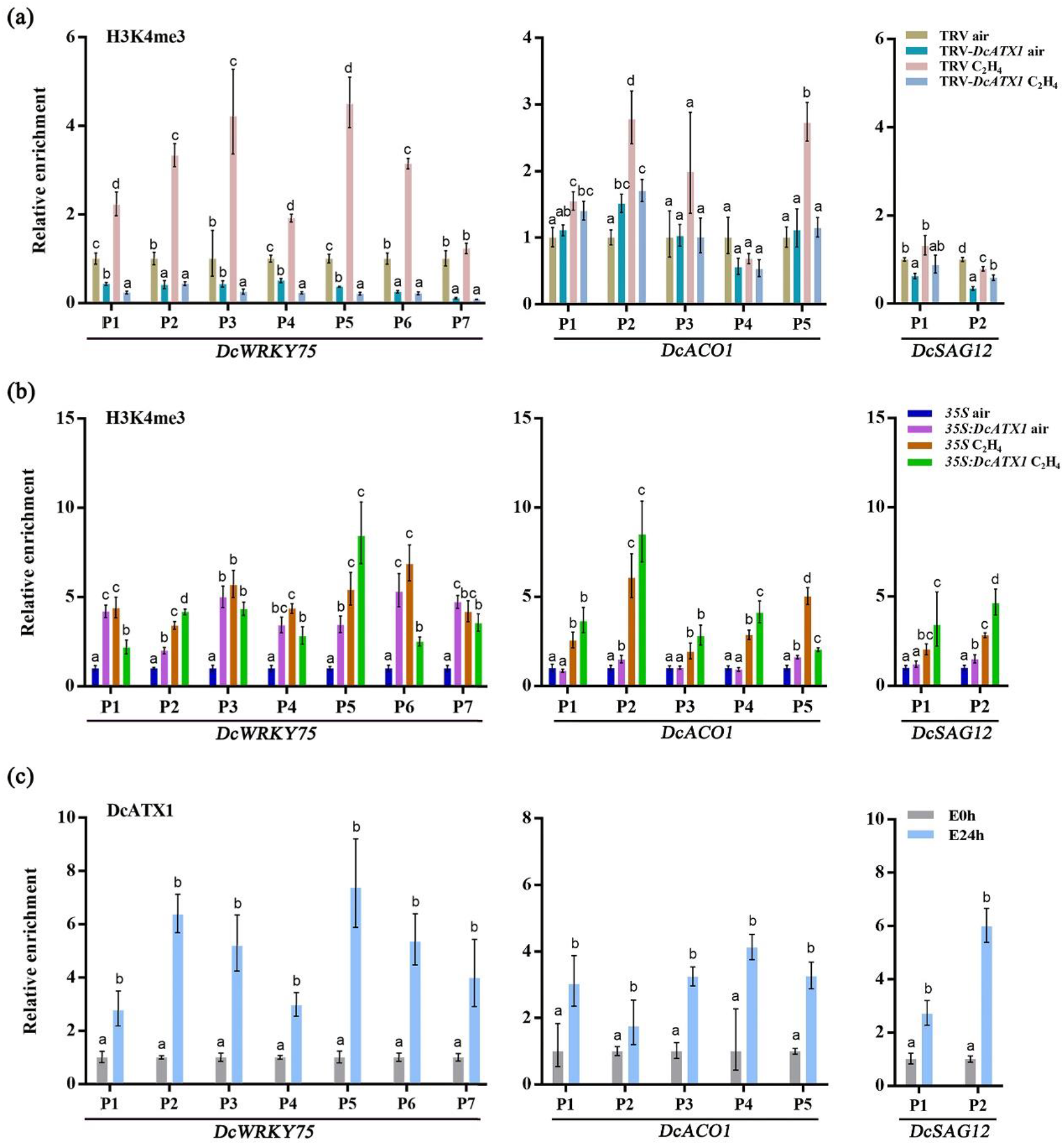
H3K4me3 level and DcATX1 accumulation at promoter regions of *DcWRKY75*, *DcACO1* and *DcSAG12* under ethylene treatment. **(a)** ChIP-qPCR detection of H3K4me3 modification at the promoters of *DcWRKY75*, *DcACO1* and *DcSAG12* in TRV control plants and TRV-*DcATX1* silenced plants without (air) and with (C_2_H_4_) ethylene treatment. **(b)** ChIP-qPCR detection of H3K4me3 modification at the promoters of *DcWRKY75*, *DcACO1* and *DcSAG12* in *35S* control plants and *35S:DcATX1* overexpression plants without (air) and with (C_2_H_4_) ethylene treatment. **(c)** DcATX1 enrichment in the promoter regions of *DcWRKY75*, *DcACO1* and *DcSAG12* without (0h) and with (24h) ethylene treatment.

We also did the ChIP-qPCR assays in *35S:DcATX1* overexpression plants and in *35S* control plants with and without ethylene treatment. We found that in the promoter of *DcWRKY75*, H3K4me3 levels were obviously increased at different regions under air condition in *35S:DcATX1* overexpression plants, and it was also increased at P2 region by ethylene treatment (Fig. 7b). For *DcACO1* promoter, H3K4me3 levels were obviously increased at P2 and P5 regions under air condition in *35S:DcATX1* overexpression plants, and it was also increased at P4 region by ethylene treatment (Fig. 7b). For *DcSAG12* promoter, H3K4me3 levels were obviously increased at P2 region no matter with or without ethylene treatment in *35S:DcATX1* overexpression plants (Fig. 7b).

These results suggested that DcATX1 promotes the transcription of *DcWRKY75*, *DcACO1* and *DcSAG12* by elevating H3K4me3 levels in their promoters.

### Ethylene promotes the accumulation of DcATX1 at the promoter regions of *DcWRKY75*, *DcACO1* and *DcSAG12*

To see whether DcATX1 regulates the H3K4me3 level by binding to the promoter regions of *DcWRKY75*, *DcACO1* and *DcSAG12*, and whether ethylene influence these binding activities, we did the ChIP-qPCR experiment in ethylene treated carnation petal.

Firstly, we conducted ChIP experiments in carnation petal without (0h) and with (24h) ethylene treatment using DcATX1 native antibody. The results clearly indicated that DcATX1 was significantly enriched in the different promoter regions of *DcWRKY75*, *DcACO1* and *DcSAG12* under ethylene treatment (Fig. 7c).

Next, we conducted ChIP experiments using transiently transformed *35S: DcATX1-GFP* carnation plants. The transiently transformed *35S: GFP* plants were used as a negative control. The results indicated that DcATX1 was also significantly enriched in the different promoter regions of *DcACO1* and *DcSAG12* under ethylene treatment (Fig. S7).

These results suggest that ethylene can promote DcATX1 accumulation at the promoter regions of *DcWRKY75*, *DcACO1* and *DcSAG12* to enhance the H3K4me3 levels, then to activate their expression to accelerate petal senescence in carnation (Fig 8).

**Figure 8.**
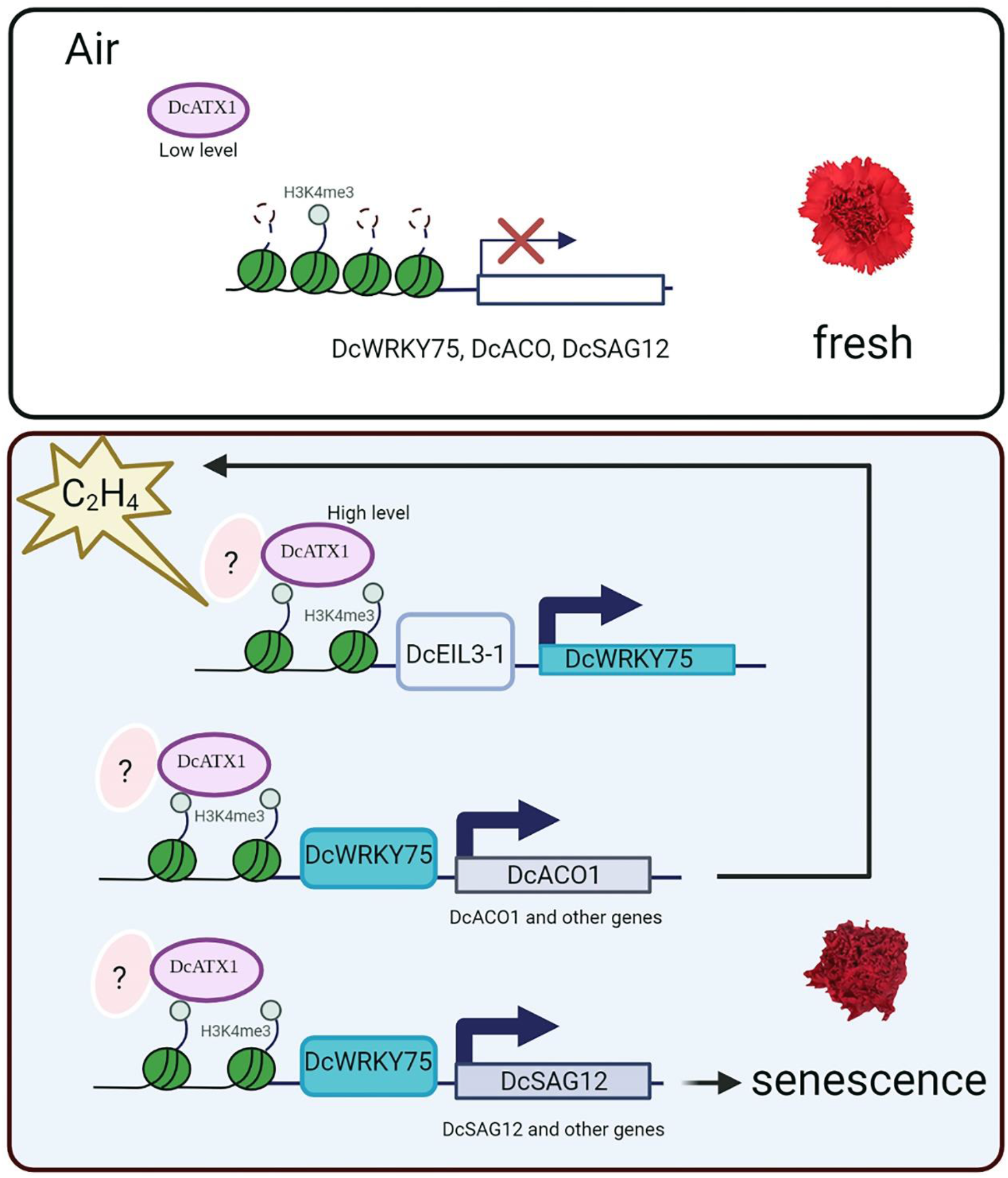
Schematic model of DcATX1 promotes ethylene induced petal senescence in carnation. In the air condition, the protein level of DcATX1 is low, so that the H3K4me3 modification on the promoters of *DcWRKY75*, *DcACO1* and *DcSAG12* is limit. In that case, the carnation flower will remain fresh; Under ethylene treatment, the DcATX1 protein will accumulate and bind to the promoters of *DcWRKY75*, *DcACO1* and *DcSAG12* by an unknown factor, so that the H3K4me3 deposition on the promoters of *DcWRKY75*, *DcACO1* and *DcSAG12* will be increased to activate their expression. The elevated expression of *DcACO1* will produce more ethylene which become into a positive feedback regulation, and the elevated expression of *DcSAG12* and other genes will promote the petal senescence, so the carnation flower will become senescent and eventually die.

## Discussion

Ethylene is important for the maturation and senescence process in plants, especially for the climacteric horticultural fruits and ornamental flowers like tomato, apple, banana, rose and carnation (Li et al., 2017; Ma et al., 2018; Liu et al., 2020; Kuang et al., 2021; Xu et al., 2021). Although numerous researches had revealed that the transcriptional regulation through transcription factors play important roles in these processes, how epigenetic regulation participated remains largely unknown (Ma et al., 2018; Chen et al., 2020; Liu et al., 2020; Tang et al., 2020).

Recently, many works have been focused on the histone methylation regulation in fruit ripening, especially for tomato fruit ripening process. The fruit ENCODE project revealed that the repressive epigenetic mark H3K27me3 plays a conserved role in regulating the expression of ethylene biosynthesis or ripening related genes during the convergent evolution of fleshy fruit ripening by analyzing 147 histone methylation profiles (Lu et al., 2018). In animals and plants, the H3K27me3 mark is mainly deposited by Polycomb Repressive Complex2 (PRC2) complexes (Margueron and Reinberg, 2011; Mozgova and Hennig, 2015). SlMSI1 (MULTICOPY SUPPRESSOR OF IRA1), a component of PRC2, has been shown to prolong the shelf life of tomato through repressing the expression of ethylene biosynthesis and fruit-ripening genes (Liu et al., 2016). Moreover, SlLHP1b (Like Heterochromatin Protein 1b), a PRC1-like protein which can interact with SlMSI1, represses fruit ripening through modulating the H3K27me3 levels in ethylene biosynthesis and ripening-related genes in tomato (Liang et al., 2020). Further, SlJMJ6, a histone lysine demethylase which can specifically demethylate H3K27 methylation, promotes tomato fruit ripening by removing H3K27 methylation of ethylene biosynthesis and ripening-regulated genes (Li et al., 2020). These studies clearly shown that the repressive epigenetic mark H3K27me3 exhibited critical function during fruit ripening in tomato and other fruits. Even though it has been shown that the active epigenetic mark H3K4me3 is essential for leaf senescence in model plant *Arabidopsis* (Ay et al., 2009; Brusslan et al., 2012; Ay et al., 2014; Brusslan et al., 2015; Liu et al., 2019; Woo et al., 2019; Li et al., 2020; Ostrowska-Mazurek et al., 2020), whether and how H3K4me3 participated in fruit ripening and flower senescence is still largely unknown. A recent story indicates that histone posttranslational modifications like H3K4me3 is crucial for fruit set in tomato (Hu et al., 2021). By using Clustered Regularly Interspaced Short Palindromic Repeats (CRISPR)/CRISPR-associated protein9 (Cas9) technique, they found that SlSDG27, a homolog of ATX1 in *Arabidopsis* which encodes a potential H3K4 methyltransferase, has potential ability to trigger the flower-to-fruit transition in tomato (Hu et al., 2021) , but the function of H3K4me3 and SlSDG27 in tomato fruit ripening is not clear. Recently, a study indicated that SlJMJ7, another histone lysine demethylase which can specifically demethylate H3K4 methylation, inhibits tomato fruit ripening by removing H3K4me3 from the promoters of ripening-regulated genes and ethylene biosynthesis genes (Ding et al., 2021), further revealed the regulation mechanism of H3K4me3 modification in fruit ripening process. But up to now, whether and how H3K4me3 participates in flower senescence is still largely undetermined.

In this work, we provide multiple lines of evidences showing that H3K4me3 plays momentous role in the ethylene induced petal senescence in carnation. Firstly, we found that the H3K4me3 levels are elevated during ethylene induced petal senescence in carnation and positively associated with expression levels of *DcWRKY75*, *DcACS1*, *DcACO1*, *DcSAG12* and *DcSAG29* (Figs. 1, 2). Secondly, we showed that DcATX1 is a H3K4 methyltransferase which can methylate H3K4 in carnation (Figs. 3, 6). Thirdly, our genetic and molecular evidence demonstrated that DcATX1 promotes ethylene induced petal senescence in carnation (Figs. 4, 5). Finally, we provided evidence showing that DcATX1 promotes the transcription of *DcWRKY75*, *DcACO1* and *DcSAG12* by targeting to their promoter regions to elevate the H3K4me3 levels (Fig. 7). Overall, this study revealed that the epigenetic modification, especially H3K4me3 which is catalyzed by DcATX1 and regulates the transcription of *DcWRKY75*, *DcACO1* and *DcSAG12*, is one of the principal molecular mechanisms to promote ethylene induced petal senescence in carnation (Fig. 8).

In the previous studies, ethylene can elevate the H3K14Ac and H3K23Ac levels to initiate the transcriptional activation (Zhang et al., 2016; Wang et al., 2017; Zhang et al., 2017; Wang et al., 2021). These give good examples that the role of plant hormone ethylene may be tightly integrated with epigenetic modification (Wang and Qiao, 2019, 2020; Wang et al., 2020). Up to now, except for histone acetylation, people do not know exactly whether and how ethylene regulates other epigenetic modifications like DNA methylation or histone methylation. Here we show that ethylene can trigger H3K4me3 accumulation on the promoter regions of *DcWRKY75*, *DcACS1*, *DcACO1*, *DcSAG12* and *DcSAG29* in carnation petal (Figs. 1, 2), which reveals a tip of the iceberg that ethylene indeed can regulate histone methylation. Future work to detect how ethylene influence H3K4me3 modification in genome-wide by Chromatin Immunoprecipitation followed by high-throughput sequencing (ChIP-seq) will be benefit for our understanding in this field.

We revealed that the gene expression of *DcATX1* was gradually decreased by ethylene treatment in carnation petal, but the protein level of DcATX1 was upregulated during this process (Fig. 3a, b). The downregulation of *DcATX1* expression maybe due to the negative feedback regulation. How ethylene promotes DcATX1 protein accumulation need to be further investigated. By ChIP-qPCR assay, we found that the enrichment of DcATX1 to the promoters of *DcWRKY75*, *DcACO1* and *DcSAG12* was significantly increased by ethylene treatment (Fig. 7c and Fig. S7), so when and how DcATX1 recognizes these sites in ethylene induced carnation petal senescence process is a momentous question. CURLY LEAF (CLF)/SDG1, another component of PRC2 which has H3K27 methylation activity, can be recruited to the chromatin by interacting with transcription factors which can bind to the polycomb response elements (Xiao et al., 2017; Zhou et al., 2018). Thus, the PRC2 recruitment relies on binding of trans-acting factors to cis-acting elements (Xiao et al., 2017; Zhou et al., 2018). Unlike CLF, the H3K27me3 demethylase REF6 can directly bind to its targets which contain a CTCTGYTY motif through its zinc finger (ZnF) domain (Cui et al., 2016; Li et al., 2016; Wang et al., 2019). The leaf senescence related H3K4me3 demethylase JMJ16 also has a ZnF domain (Liu et al., 2019). Genetic and molecular evidence shown that the ZnF domain of JMJ16 is not required for binding to its targets like *WRKY53* and also not required for H3K4me3 enrichment (Liu et al., 2019). This means that different histone methylation enzymes using different strategies to recognize their targets. Since DcATX1 can bind to the promoter of *DcWRKY75* under ethylene treatment (Fig. 7c) and *DcWRKY75* is a direct target gene of DcEIL3-1 (Xu et al., 2021), we detect the interaction of DcATX1 with DcEIL3-1 by yeast two-hybrid (Y2H) assay. There is no interaction between DcATX1 and DcEIL3-1 (Fig. S8). Further, since DcATX1 also binds to the promoters of *DcACO1* and *DcSAG12* under ethylene treatment (Fig. 7c and Fig. S7) and *DcACO1* and *DcSAG12* are the direct target genes of DcWRKY75 (Xu et al., 2021), we also detect the interaction of DcATX1 with DcWRKY75 by Y2H assay. However, there is also no interaction between DcATX1 and DcWRKY75 (Fig. S8). These results indicated that DcATX1 is not directly recruited by DcEIL3-1 or DcWRKY75 under ethylene treatment. Using omics method like ChIP-seq to search the DcATX1 binding sites or binding motifs *in vivo* by DcATX1 native antibody and by using immunoprecipitation combined with mass spectrometry (IP/MS) to detect protein interactors of DcATX1 is important for revealing its regulation mechanism in carnation petal senescence in the future. Moreover, due to the limitations of VIGS gene silencing approach, the TRV-*DcATX1* silenced plants still show some response to ethylene, further work in constructing carnation *dcatx1* true mutant by CRISPR/Cas9 technique to verify its function in regulating petal senescence is necessary. By the way, whether and how other H3K4 methylation related SDG proteins and JMJ proteins involved and participate in ethylene induced petal senescence in carnation is need to be further investigated.

In conclusion, we revealed the function of a H3K4 methyltransferase DcATX1 in ethylene induced petal senescence in carnation, which gave us a new view about the involvement of epigenetic modification in flower senescence. Ethylene can promote DcATX1 binding to the promoter regions of *DcWRKY75*, *DcACO1* and *DcSAG12* to elevate the active H3K4me3 mark to enhance their expression, thus accelerating petal senescence in carnation (Fig 8). This study reveals the tip of the iceberg of how epigenetic regulation participates in the transcriptional regulation network in petal senescence. Manipulation of *DcATX1* homolog genes by gene editing technique maybe benefit to cultivate longer vase life of ethylene insensitive cut flowers.

## Materials and methods

### Plant materials and growth conditions

Cut carnation (‘Master’) flowers used in this study were collected from a commercial grower (Kunming, China) and transported to the laboratory within 24h. Stems were recut to 35 cm in length and rehydrated in deionized water (DW) and then held in refreshed DW. Flower opening stages were recorded as previously described (Kong et al., 2017). The *Arabidopsis* Columbia-0 (Col-0) were grown in an artificial growth chamber at 22℃ under long days (16h light/8h dark cycle) until use.

### Ethylene treatments

FBS flowers were sealed in 10 liters airtight chambers with 10 ppm ethylene at 25°C for different times. Flowers exposed to air were used as the control. 1 mol/L NaOH was placed in the chamber to prevent the accumulation of CO_2_. After treatment, the samples used for the gene expression analysis were collected and immediately frozen in liquid nitrogen, then stored at -80℃.

### Plasmid construction and plant transformation

Silencing of *DcATX1* in carnation flowers by VIGS technique was performed as previously described (Cheng et al., 2018; Xu et al., 2021). For construction of the *DcATX1* VIGS vector, a 320 bp fragment from the 5’ end of the *DcATX1* coding region was amplified and cloned into the VIGS vector pTRV2. To construct pCAMBIA1300-*DcATX1* vector, the full-length genomic coding region of *DcATX1* was amplified using gene-specific primers and then inserted into pCAMBIA1300 vector with a C-terminal GFP tag. For transient transformation of carnation, the pCAMBIA1300-*DcATX1* construct, pTRV2-*DcATX1* construct as well as pTRV1 and pTRV2 were transformed into *Agrobacterium tumefaciens* cells (strain GV3101) and then cultured in Luria-Bertani medium for 12 h. The cultures were resuspended in infiltration buffer (10 mM MgCl_2_, 200 mM Acetosyringone, 10 mM MES) to a final OD_600_ of approximately 1.5. pTRV2 were used as control. After incubating for 4 hours in the dark at 28℃, the carnation flowers or 0.6mm diameter discs excised from the carnation petals were immersed in the pCAMBIA1300-*DcATX1* and pCAMBIA1300 bacterial suspension for transient overexpression of *DcATX1* or in the mixtures of bacterial suspension containing an equal ratio (v/v) of pTRV1 and pTRV2 or pTRV1 and pTRV-*DcAX1* for VIGS assay, followed by infiltrating under a vacuum at 0.7 MPa. After vacuum infiltration, the petal discs or carnation flowers were washed and placed in sterile water in the dark at 7-8°C for 3 d, followed by keeping at 23°C until sampling. At least 16 petal discs were used for each treatment, and three replicates were performed for each treatment. For VIGS of carnation plants, around 4 plants were used in each of five independent experiments. The pCAMBIA1300-*DcATX1* was stably transformed into *Arabidopsis* Col-0 WT using the floral dip method (Clough and Bent, 1998).

### Measurement of electrolyte leakage rates

Electrolyte leakage rates were measured as described previously (Wu et al., 2017). Briefly, fifteen carnation petal discs from each treatment were immersed in 15 ml of 0.4 M mannitol and shaken for 3h at room temperature. The initial conductivity of the solution was measured with a conductivity meter (ST3100C) followed by determination of total conductivity after the sample was incubated at 85°C for 20 min. The electrolyte leakage rates were calculated as the percentage of initial conductivity to the total conductivity.

### Subcellular localization analysis

The subcellular localization of DcATX1 was carried out by infiltrating the *Nicotiana benthamiana* leaves with *Agrobacterium tumefaciens* (GV3101) which carrying the recombinant vector pCAMBIA1300-*DcATX1*. The nucleolus marker gene fused with mCherry-RFP was used as nucleus maker (Xu et al., 2021). After infiltrating for 3 d, the infiltrated leaves were imaged in tobacco leaf epidermal cells using a laser confocal microscope (Leica TCS SP8).

### RT-qPCR

Total RNA samples were extracted using the TRlzol ®Reagent (Invitrogen) and 1 µg of total RNA was converted into cDNA using HiScript 1st Strand cDNA Synthesis Kit (Vazyme) following the manufacturer’s instructions. RT-qPCR reactions were performed on BIO-RAD CFX Connect real time system using a HieffTMqPCR SYBR& Green Master Mix (Yeasen) (Zhang et al., 2014; Zhang et al., 2016). The transcript levels of genes were normalized to the internal control gene *DcUbq3-7* following the 2^-ΔΔCt^ method (Nomura et al., 2012).

### Protein extraction and western blot

For total protein extraction, frozen carnation flower samples were ground in liquid N_2_ and extracted with extraction buffer (100 mM Tri-HCl, pH 7.5, 100 mM NaCl, 5 mM EDTA, 10 mM N-ethylmaleimide, 5 mM DTT, 10 mM β-mercaptoethanol, 1% SDS), and centrifuged at 13,000g for 3 min at 4℃. The supernatant was collected and prepared for western blot analysis. Western blot analysis were performed as previously described (Zhang et al., 2016; Zhang et al., 2018). The relative amounts of protein levels were calculated by ImageJ (https://imagej.nih.gov/ij/). Antibodies used in western blot were anti-H3K4me3 (Abclonal A2357, 1:2000 dilution), anti-H3K4me2 (Abclonal A2356, 1:2000 dilution), anti-H3K4me1 (Abclonal A2355, 1:2000 dilution), anti-H3 (Abclonal A2348, 1:2000 dilution), anti-Rubisco (Abkine A01110, 1:2000 dilution), anti-MBP (Abclonal AE016, 1:2000 dilution) and anti-DcATX1 (Abclonal, 1:500 dilution).

### *In vitro* histone methyltransferase assay

An *in vitro* histone methyltransferase assay was performed as previously described (Guo et al., 2010). The 3′ region of the *DcATX1* cDNA encoding an SET domain was amplified using specific PCR primers and cloned into the pMAL-C2 vector. MBP-DcATX1C fusion protein was expressed in *E. coli* BL21 (DE3) and purified using Amylose Resin (MBP, New England Biolabs) according to the manufacturer’s introduction as well as recombinant *Arabidopsis* histone MBP-H3_1-57_. After purification of MBP-fused proteins, histone methyltransferase assay was carried out. Briefly, 50µl reaction mixtures containing substrate (MBP-H3_1-57_), enzyme (MBP-DcATX1C) and S-adenosyl-L-methionine (SAM; NEB) were incubated for 0h, 1h, 2h, 4h, 8h, 12h and 24h at 37°C. Reactions were stopped by boiling in sodium dodecyl sulfate (SDS) loading buffer. After the histone methyltransferase assay, the reaction mix was separated by SDS/PAGE gel electrophoresis, dried, and exposed to films, as well as analyzed by western blot analysis using specific antibodies.

### ChIP-qPCR

ChIP-qPCR was performed according to a previously published protocol (Zhang et al., 2016; Zhang et al., 2017). Briefly, chromatin isolated from carnation petals were fixed in 1% formaldehyde and sonicated into DNA fragments. 6% of the sonicated chromatin was saved as the input. The left samples were subsequently incubated with 2 µl of antibodies of anti-H3K4me3 (Abclonal A2357), anti-GFP (Abkine A02020) and Magnetic Protein G Beads (Promega, G747A) overnight at 4°C with gentle agitation. The input and eluted DNA were amplified by quantitative real-time PCR to determine the enrichment of DNA immunoprecipitated.

### Y2H assay

The full-length cDNA sequences of *DcATX1*, *DcWRKY75* and *DcEIL3-1* were cloned into the yeast vector pGBKT7 and pGADT7 separately to examine the interactions between DcATX1, DcWRKY75 and DcEIL3-1. The Y2H Gold yeast strain (Clontech) was transformed with appropriate pGADT7 and pGBKT7 constructs. Yeast strains were grown on synthetic dropout (SD) medium minus Trp and Leu (SD/-Trp-Leu) plates for 3 days at 30°C, and then were spotted on the selective plates of SD/-Trp-Leu-His-Ade. All primers used in this study are listed Data S2.

### Statistical analysis

All experiments were performed with at least three biological replicates. Asterisks indicate significant differences (Welch’s t-test, * P < 0.05, **P < 0.01, ***P < 0.001). Different letters indicate significant differences by using Welch’s t test comparisons (P <0.05). Error bars represent ±SD.

## Accession numbers

Sequence data from this article can be found in the TAIR website (https://www.arabidopsis.org) under the following AGI codes: ATX1/SDG27 (AT2G31650), ATX2/SDG30 (AT1G05830), ATX3/SDG14 (AT3G61740), ATX4/SDG16 (AT4G27910), ATX5/SDG29 (AT5G53430), ATXR3/SDG2 (AT4G15180), ATXR7/SDG25 (AT5G42400), ASHH1/SDG26 (AT1G76710), ASHH2/SDG8 (AT1G77300), ASHR3/SDG4 (AT4G30860), SAG12 (AT5G45890), SAG29 (AT5G13170) and ACTIN2 (AT3G18780).

## Acknowledgements

We thank Prof. Rongcheng Lin (IBCAS, China) for his critical suggestion. This work was supported by grants from the Fundamental Research Funds for the Central Universities (2662019PY049), by grants from Thousand Youth Talents Plan Project and by Start-up Funding from Huazhong Agricultural University to F.Z.

## Data availability

The data supporting the findings of this study are available within the paper and its Supplementary information files.

## Supplementary information

**Figure S1.** The expression levels of *ATX1*, *ATX2*, *ATX3*, *ATX4*, *ATX5*, *ATXR7*, *ASHH1*, *ASHH2*, *ASHR3* shown by eFP browser.

**Figure S2.** Phylogenetic tree and expression profiles of H3K4 methylation related SDG proteins.

**Figure S3.** Proteins sequence alignment and domain analysis of DcATX1.

**Figure S4.** *DcATX1* silencing delays senescence in carnation petal discs.

**Figure S5.** *DcATX1* overexpression accelerates senescence in carnation petal discs.

**Figure S6.** Overexpression of *DcATX1* promotes flower senescence in *Arabidopsis*.

**Figure S7.** DcATX1 enrichment in the promoter regions of *DcACO1* and *DcSAG12* without (air) and with (C_2_H_4_) ethylene treatment.

**Figure S8.** DcATX1 do not interact with DcEIL3-1 and DcWRKY75.

**Data S1.** FPKM values of H3K4 methylation related SDG genes in ethylene treated carnation petal transcriptome.

**Data S2.** Sequences of primers used in this study.

